# A yeast-based tool for mammalian DGATs inhibitors screening

**DOI:** 10.1101/2022.07.06.498986

**Authors:** Peter Gajdoš, Rodrigo Ledesma-Amaro, Jean-Marc Nicaud, Tristan Rossignol

## Abstract

Dysregulation of lipid metabolism is associated with obesity, metabolic diseases but there is also increasing evidences of a relationship between lipid bodies (LBs) excess and some cancer. LBs synthesis requires diacylglycerol acyltransferases (DGATs) which catalyses the last step of triacylglycerol (TAG) synthesis, the main storage lipid form in lipid bodies. DGATs and in particular DGAT2, are therefore considered as potential therapeutic targets for the control of these pathologies. Here, the murine and the human DGAT2 were overexpressed in the oleaginous yeast *Yarrowia lipolytica* deleted for all DGATs activities, for evaluating the functionality of the enzymes in this heterologous host and to evaluate DGAT activity inhibitors. This work provide evidence that mammalian DGATs expressed in *Y. lipolytica* is a useful tool for screening chemical libraries to identify potential inhibitors or activators of these enzymes of therapeutic interest.

## Introduction

The diacylglycerol acyltransferase (DGAT) enzymes catalyse the final committed step of the triacylglycerol (TAG) biosynthesis by esterification of a fatty acyl moiety to a diacylglycerol. These neutral lipids are stored in organelles called lipid bodies (LBs) in mammalian adipocytes tissues but also in most eukaryotic cells and in some prokaryotes, as energy molecules or membrane synthesis synthesis reservoir. In eukaryotes, TAGs are mainly synthetized by DGAT1 and DGAT2, belonging to two different gene families. DGAT1 and DGAT2 have different roles in TAG synthesis in humans. DGAT1 is highly expressed in the small intestine and has a role in fat absorption while DGAT2 is expressed in liver and adipose tissues and is responsible for endogenous synthesis of TAG (Cases et al. 1998; Cases et al. 2001). *dgat1* knockout mice are viable with minor impact on TAG levels and are resistant to diet-induced obesity (Smith et al. 2000). In contrast, *dgat2* knockout mice present severe TAG decrease and die shortly after birth (Stone et al. 2004). TAGs excess in tissues is a hallmark of obesity. Therefore, DGATs are considered as potential therapeutic inhibition targets for the control of obesity, but also for some diseases related to lipid absorption in the intestine. Moreover, recent studies revealed that high levels of LBs are also associated with breast cancer (Nisticò et al. 2021) as well as with higher tumor aggressiveness and chemotherapy resistance (Tirinato et al. 2017). Interestingly DGAT2 is constitutively activated in various cancers including breast cancer (Hernández-Corbacho and Obeid 2019). Accordingly, Nistico et al. observed that the combined effect of a DGAT2 inhibitor pre-treatment (PF-06424439) and radiation enhanced radiosensitivity of MCF7 breast cancer cells (Nisticò et al. 2021). In addition, importance of DGAT2-mediated regulation of TAG metabolism in triple negative breast cancer has been recently highlighted (Almanza et al. 2022). Therefore, DGAT2 appears as a new potential therapeutic target in the treatment of breast cancer (Hernández-Corbacho and Obeid 2019). Because of the lethality of *dgat2* knockout mice model, specific inhibitors that tightly controlled inhibition of DGAT2 are required. Compound libraries targeting obesity as well as cancer should be evaluated in a system that could mimic human and mouse DGAT2 structure and activity with ease and high throughput screening capacity.

Being able to express these enzymes in a simple heterologous model would provide an efficient and versatile tool to characterize these enzymes and potential inhibitors. To do so, in this work, the oleaginous yeast *Yarrowia lipolytica* has been used as a heterologous host. This yeast is particularly valuable in this context. It has been a model for lipid metabolism for decades and has a high enzyme production capacity (Nicaud 2012), it can produce large LBs and it is easy to manipulate thanks to the numerous modern genetic engineering toolBox now available (Larroude et al. 2018). In particular, a strain deleted for all the genes coding for enzymes with DGAT activities (Q4) is available (Beopoulos et al. 2012). This strain is not able to form LBs anymore. Previous work has shown that DGAT activity in *Y. lipolytica* can be easily validated, characterized and modulated by overexpression approaches in this genetic background, allowing restoration of LBs formation and TAGs accumulation (Aymé et al. 2015; Gajdoš et al. 2016; Gajdoš et al. 2019). Those previous works established the efficiency and versatility of the heterologous expression of DGAT in this particular host.

In order to determine whether the heterologous constructs could potentially be used as tools to measure the activity of these enzymes and thus useful for screening chemical libraries to identify regulatory molecules, the murine and the human DGAT2 were overexpressed in the above mentioned Q4 strain, as well as the oleaginous fungus *Umbelopsis rhamaniana* DGAT2, the first DGAT2 identified and expressed in heterologous host (Lardizabal et al. 2001), and the *Y. lipolytica* DGATs for comparison. MmDGAT2 and HsDGAT2 have already been expressed in heterologous systems including insect cells and yeast, but mainly for *in vitro* activity assays (Cases et al. 2001; Turkish et al. 2005; Yen et al. 2005; Stone et al. 2006; Kim et al. 2014). DGATs overexpression in the Q4 chassis strains were therefore first evaluated for the LBs restoration phenotype and TAG accumulation to evaluate the capacity to use them as *in vivo* screening tools for DGATs inhibitors candidate drugs. These strains were then exposed to known inhibitory molecules of mammalian DGAT1 and DGAT2. The results showed that the DGATs are functional in our chassis and that inhibitors conserved their specificities and efficacy. This work provided proof of principle for using these strains as a screening system for libraries of molecules to discover new inhibitors or activators of these enzymes of particular therapeutic interest.

## Results and discussion

### LBs forming phenotype complementation

The Q4 chassis strain used in this study is unable to form LBs. Therefore, heterologous DGAT functionality can be easily evaluated by simple observation of LBs restoration directly supporting enzymatic activity in the heterologous host. The *Y. lipolytica* YlDGAT1 and YlDGAT2 overexpressed in the Q4 chassis were previously validated (Beopoulos et al. 2012; Gajdoš et al. 2016) and serve as controls for the Q4 strains overexpressing MmDGAT2, UrDGAT2 and HsDGAT2. All strains show a complementation phenotype by formation of LBs (Fig. 1). YlDGAT2 overexpression being the highest, as expected as it is the main DGAT for lipid accumulation in LBs in *Y. lipolytica* (Gajdoš et al. 2016) while HsDGAT2 being the lowest in the condition tested with small LBs formation.

**Fig. 1.**
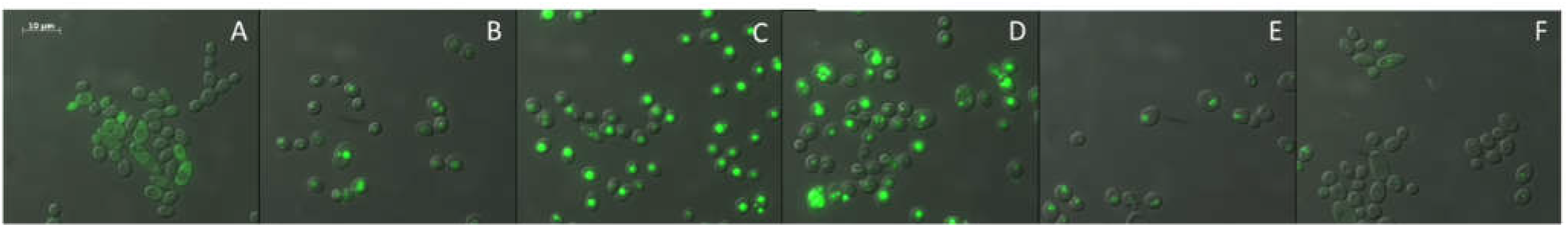
Phenotype complementation of strains overexpressing DGAT. A) Strain Y1880 Q4. B) Strain Y1884 Q4+YlDGAT1. C) Strain Y1892 Q4+YlDGAT2. D) Strain Y3137 Q4+MmDGAT2. E) Strain Y4592 Q4+UrDGAT2. F) Strain Y7378 Q4+HsDGAT2.

### DGAT2 Inhibitor activity on HsDGAT2 evaluation

As the HsDGAT2 is functional in *Y. lipolytica*, we therefore evaluated FP-06424439, a specific inhibitor of DGAT2, for its capacity to inhibit the LBs formation complementation phenotype in yeast. Serial concentration of FP-06424439 were tested against the strain Y7378 overexpressing the Human DGAT2. In these experiments, we increase the C/N ratio in the medium to 30 as it increases the TAG accumulation in *Y. lipolytica* (Gajdoš et al. 2016) and will consequently improve LBs visualization. Accordingly, LBs appear bigger in the Y7378 strain without treatment (Fig. 2). Small inhibition of LBs formation can be observed at 12.5 µg/ml and increases at 25 µg/ml, where almost no LBs can be observed. At 50 µg/ml, no LBs are formed (Fig. 2). The different concentration of PF-06424439 have no impact on the fitness of the strain as growth is no affected for all the concentrations tested (Fig. 3).

**Fig. 2.**
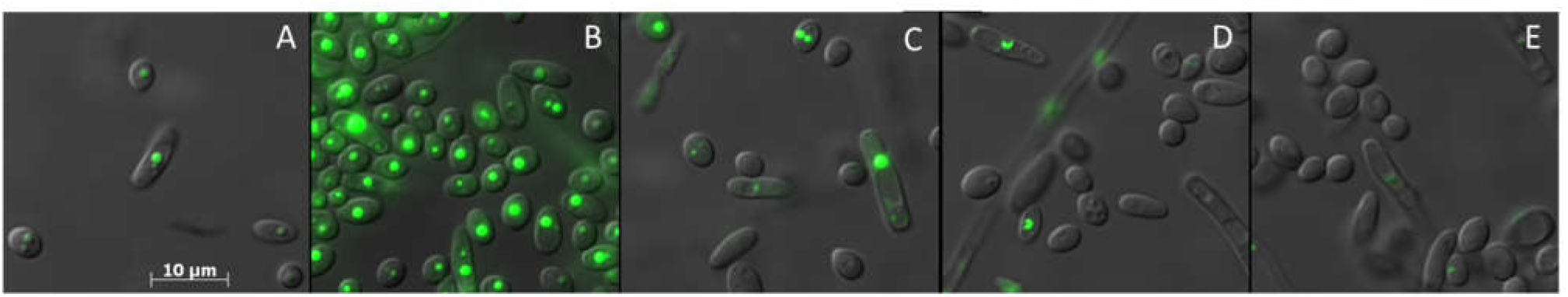
LBs formation inhibition evaluation in Y7378 Q4+HsDGAT2 using different concentration of PF-06424439. A) 0 µg/mL., B) 6.25 µg/mL., C) 12.5 µg/mL., D) 25 µg/mL., E) 50 µg/mL.

**Fig. 3.**
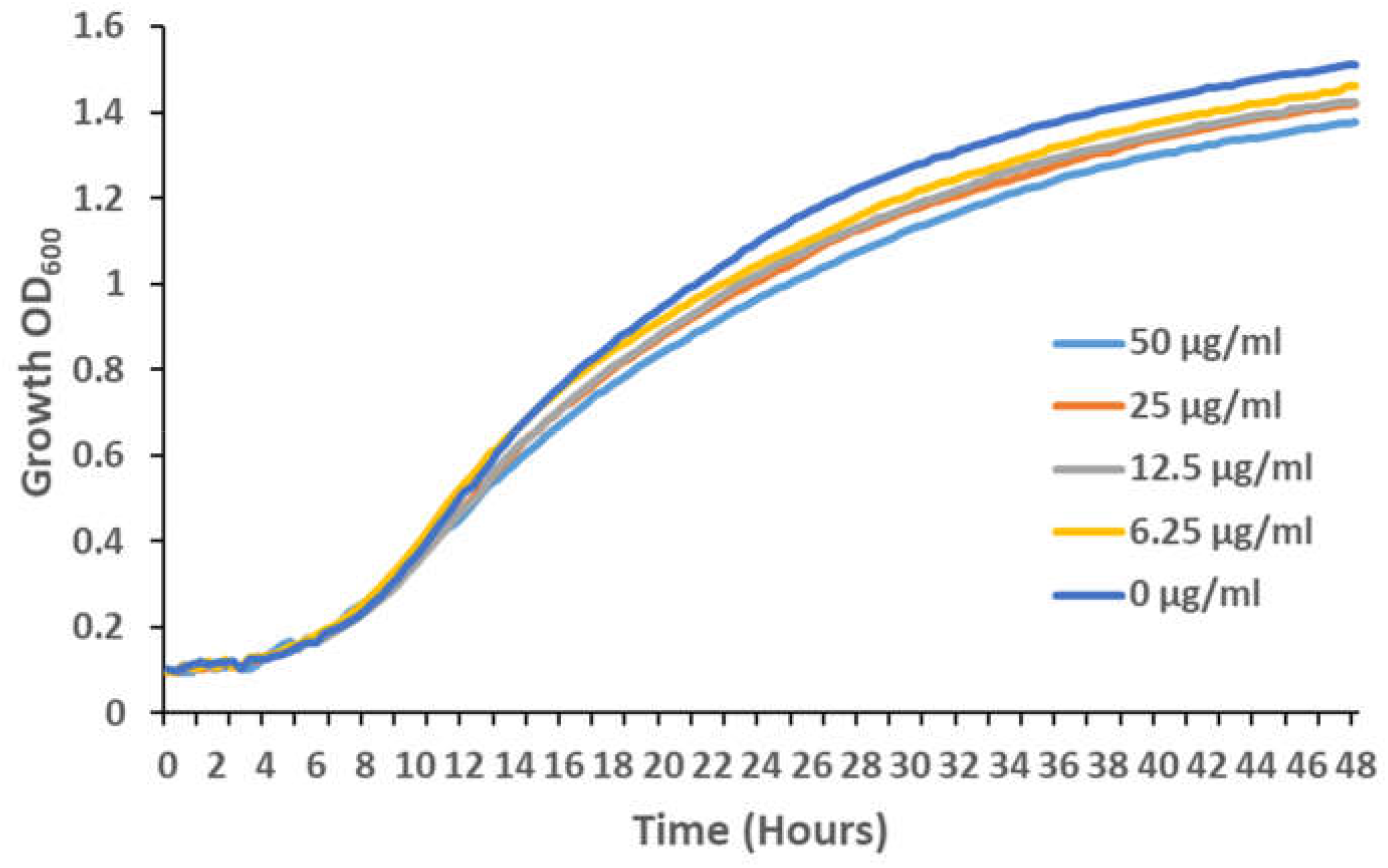
Y7378 (Q4+HsDGAT2) growth curve on microtiterplate with increasing concentration of PF-06424439.

### Specificity of DGAT inhibitors on DGATs from different origins

The previous experiment established a minimal concentration that inhibits LBs formation without affecting growth for the DGAT2 specific inhibitor PF-06424439. This is also in the range of concentrations for which LBs are reduced in MCF7 breast cancer without affecting cells viability (Nisticò et al. 2021). Therefore, we tested this inhibitor at the same concentration on the six DGAT overexpressing strains selected in this study to evaluate the DGAT specificity. PF-046020110, a DGAT1 specific inhibitor, was also evaluated as a control. PF-046020110 has no effect on the different strains overexpressing DGAT even YlDGAT1 at the concentration tested (Fig. 4). PF-06424439 inhibit LBs formation of the strain overexpressing the HsDGAT2 as demonstrated in Fig. 2, and has a similar effect on MmDGAT2, while no inhibition was observed for the other strains overexpressing *Y. lipolytica* DGATs or *U. rhamaniana* DGAT2, indicating a specificity for mammalian DGAT2 (Fig. 4). No growth defects were observed for all strains with any of the compound (supplementary data).

**Fig. 4.**
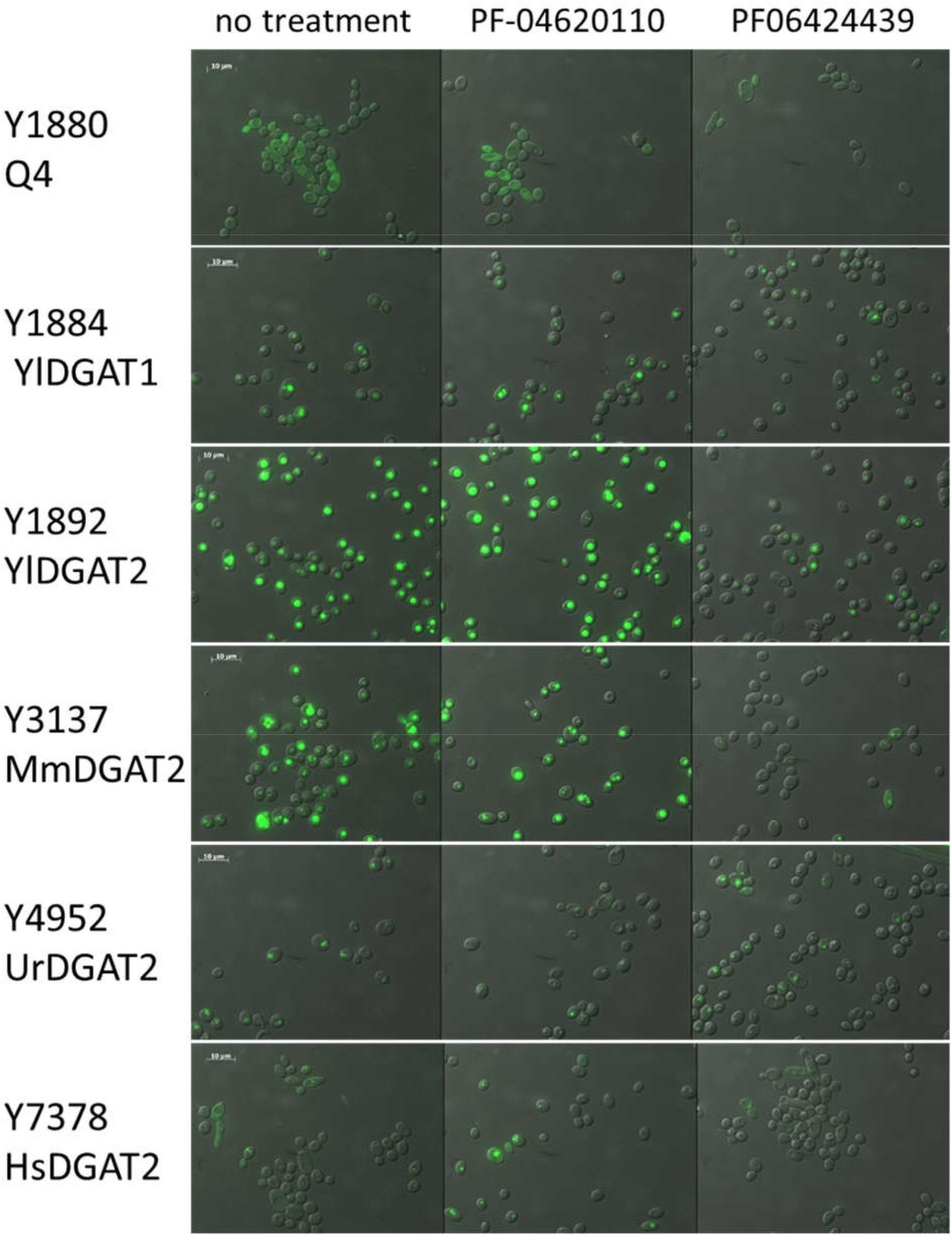
LBs formation inhibition evaluation in the strains overexpressing different DGATs using PF-06424439 and PF-046020110 as inhibitors at 25µg/mL.

## Conclusion

Here, we provide strong arguments that *Y. lipolytica* can serve as an efficient platform for the expression and study of heterologous DGATs and for the screening of therapeutic molecules targeting these enzymes. The mammalian heterologous DGATs cloned here are from cDNA libraries and not optimized for expression in *Y. lipolytica*. Nevertheless, we show that the variants used are functional and the sequences are sufficiently conserved to retain activity in the heterologous host while keeping the drug inhibitor specificity. Thus, this strain platform is perfectly suited to screen compounds libraries as mammalian DGAT2s react as in their natural environment. In addition, the proof of principle presented here is performed in microtiterplate, which is well adapted to high throughput screening thanks to the simple phenotype evaluation.

## Material and methods

### Compounds, media and culture conditions

PF-06424439 and DMSO for PF-046020110 were obtained from Sigma-Aldrich. Stock solutions were prepapred by resuspending the powder at 5mg/mL in sterile distilled water for PF-06424439 and in pure DMSO for PF-046020110. *E. coli* strains were grown in Luria-Bertani broth medium complemented with 50 μg/mL kanamycin or 100 μg/mL ampicillin when required. For yeast growth and transformant selection, minimal YNB medium, composed of 0.17% (w/v) yeast nitrogen base (without amino acids and ammonium sulfate), 0.5% (w/v) NH_4_Cl, 50 mM phosphate buffer (pH 6.8), and 2% (w/v) glucose was used. Leucine was added at a final concentration of 0.1 g/L when required. For higher lipid accumulation experiments, similar YNB medium with 0.15% (w/v) NH_4_Cl and 3% (w/v) glucose was used, which corresponded to a carbon-to-nitrogen ratio of 30 (C/N 30). Solid media were complemented with 1.6% agar.

For 96 wells microtiterplates, yeasts were pre-cultured in YNB overnight at 28°C, washed and diluted in fresh YNB medium at an optical density at 600 nm of 0.2. 100 µl of this dilution was mixed with 100 µl of inhibitor solution diluted in YNB at the required concentration. Cultures were grown at 28◦C under constant agitation on a Biotek Synergy MX microtiterplate reader (Biotek Instruments) and monitored by measuring optical density at 600 nm every 20 min for 72 h.

### Plasmid and strains construction

*UrDGAT2* (GenBank accession number: AAK84179.1) was synthetized and codon optimized by Genscript. The gene was cloned under the pTEF promoter between BamHI and AvrII cloning sites in the overexpression JMP62 vector (Nicaud et al. 2002) containing the selective marker *URA3*, to generate JMP2881 plasmid. *MmDGAT2* were PCR-amplified from cDNA cloned vectors (Yen et al. 2005) using primers that allowed Gateway cloning by introducing attb sequences (Attb1-DGAT2_forward GGGGACAAGTTTGTACAAAAAAGCAGGCTATGAAGACCCTCATCGCCGCCTACTCCGGG; Attb2-DGAT2_reverse GGGGACCACTTTGTACAAGAAAGCTGGGTCTCAGTTCACCTCCAGCACCTCAGTCTCTG). PCR fragments were cloned into the Gateway® vector pDONR207 (Invitrogen) using Gateway BP clonase (Thermo Fisher Scientific) to generate plasmid JMP1783 and transferred into the *Y. lipolytica* Gateway expression vector JMP1529 (Leplat et al. 2015) using Gateway LR clonase (Thermo Fisher Scientific) giving rise to the plasmids JMP1785 (*pTEF-MmDGAT2-URA3ex*). The *HsDGAT2* cDNA clone (NM_032564) was bought from Genscript and was amplified with the same primers as for *MmDGAT2* (one nucleotide difference and no change in the amino acid sequences) to remove the C-terminal tag present in the vector and introducing the attb sequences for Gateway® cloning. PCR fragments was cloned into the Gateway® vector pDONR207 (Invitrogen) to generate plasmid JME4451 and transferred into the *Y. lipolytica* Gateway expression vector JMP1529 (Leplat et al. 2015), giving rise to the plasmids JMP4468 (*pTEF-HsDGAT2-URA3ex*). The nucleoSpin plasmid easypure Kit (Macherey-Nagel) was used for plasmid purification and expression cassettes were sequence verified. Expression cassettes from the *NotI*-digested plasmids JMP2881, JMP1785 and JMP4468 were used for *Y. lipolytica* transformation in the Y1877 strain (the Q4 strain) which lacked the four acyltransferases (Beopoulos et al. 2012) using the lithium acetate method (Le Dall et al. 1994), creating strains Y4952 (Q4-UrDGAT2), Y3137 (Q4-MmDGAT2) and Y7378 (Q4-HsDGAT2) respectively. Strains Y1880 (Q4), strains Y1884 (Q4-YlDGAT1) and Y1892 (Q4-YlDGAT2) were described previously (Beopoulos et al. 2012). All the plasmids and strains used in this study are listed in table 1 and 2, respectively.

**Table 1.** List of plasmids used in this study

| Plasmids | genotype | References |
| --- | --- | --- |
| JMP1529 | Gateway expression vector | (Leplat et al. 2015) |
| JMP1046 | Expression vector | (Nicaud et al. 2002) |
| JMP2881 | JME1046+UrDGAT2 | This work |
| JMP1783 | pDONR207+MmDGAT2 | This work |
| JMP1785 | JMP1529+MmDGAT2 | This work |
| JMP4451 | pDONR207-HsDGAT2 | This work |
| JMP4468 | JME1529-HsDGAT2 | This work |

**Table 2.** List of *Y. lipolytica* strains used in this study

| Strains | Genotype | References |
| --- | --- | --- |
| Y1877 (Q4) | *leu2-270 ura3-302 Δdga1Δlro1Δare1Δdga2* | (Beopoulos et al. 2012) |
| Y1880 | *leu2-270 ura3-302 Δdga1Δlro1Δare1 ::URA3 Δdga2* | (Beopoulos et al. 2012) |
| Y1884 | *leu2-270 ura3-302 Δdga1Δlro1Δare1Δdga2 pTEF-YIDGAT1-URA3ex* | (Beopoulos et al. 2012) |
| Y1892 | *leu2-270 ura3-302 Δdga1Δlro1Δare1Δdga2 pTEF-YIDGAT2-URA3ex* | (Beopoulos et al. 2012) |
| Y3137 | *leu2-270 ura3-302 Δdga1Δlro1Δare1Δdga2 pTEF-MmDGAT2-URA3ex* | This work |
| Y4952 | *leu2-270 ura3-302 Δdga1Δlro1Δare1Δdga2 pTEF-UrDGAT2-URA3ex* | This work |
| Y7378 | *leu2-270 ura3-302 Δdga1Δlro1Δare1Δdga2 pTEF-HsDGAT2-URA3ex* | This work |

### Fluorescence microscopy

For LBs staining, cells were stained at room temperature by a 10-min incubation with BODIPY 493/503 (Invitrogen) at 1 μg/mL. Images were taken using a Zeiss Axio Imager M2 microscope with a 100X oil immersion objective, equipped with an HXP 120 C lamp.

## Acknowledgments

We would like to thank Joel Haas for kindly providing us the *MmDGAT2* vectors.

## Notes

### Competing Interest Statement

The authors have declared no competing interest.

