## Supplementary figures and images for "A yeast-based tool for mammalian DGATs inhibitors screening"

### supplementary data

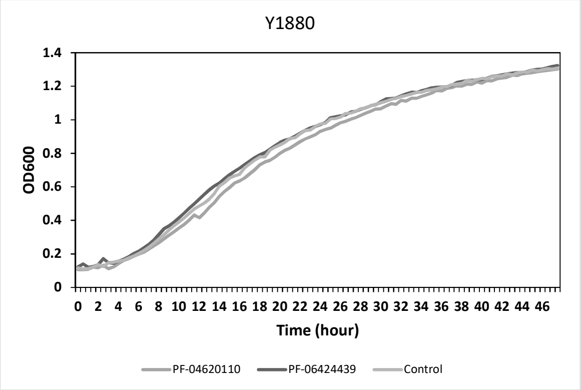

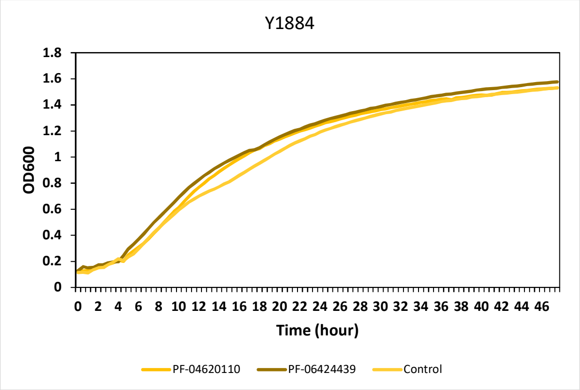

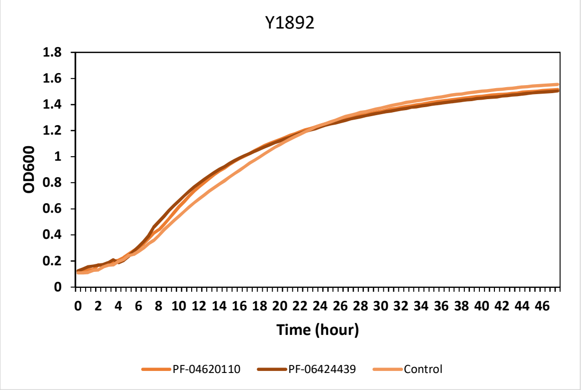

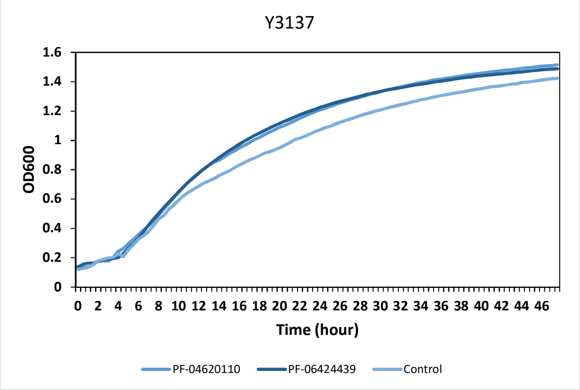

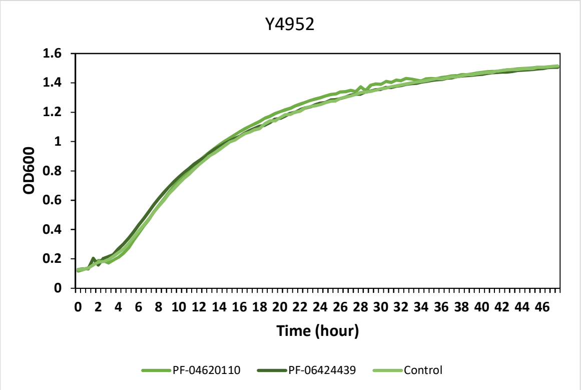

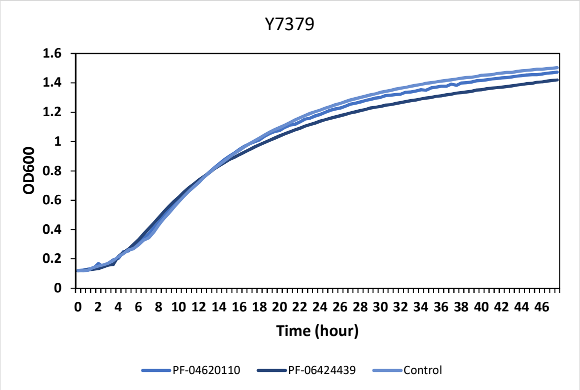
